## Supplementary Information for "Ordering and topological defects in social wasps’ nests"

### Ordering and topological defects in social wasps' nests
### Supplementary Information


Shivani Krishna[1,*], Apoorva Gopinath[1] and Somendra M. Bhattacharjee[2]

[1]Department of Biology, Ashoka University, Sonepat 131029, India
[2]Department of Physics, Ashoka University, Sonepat 131029, India


#### S1. Plots of pair correlation functions

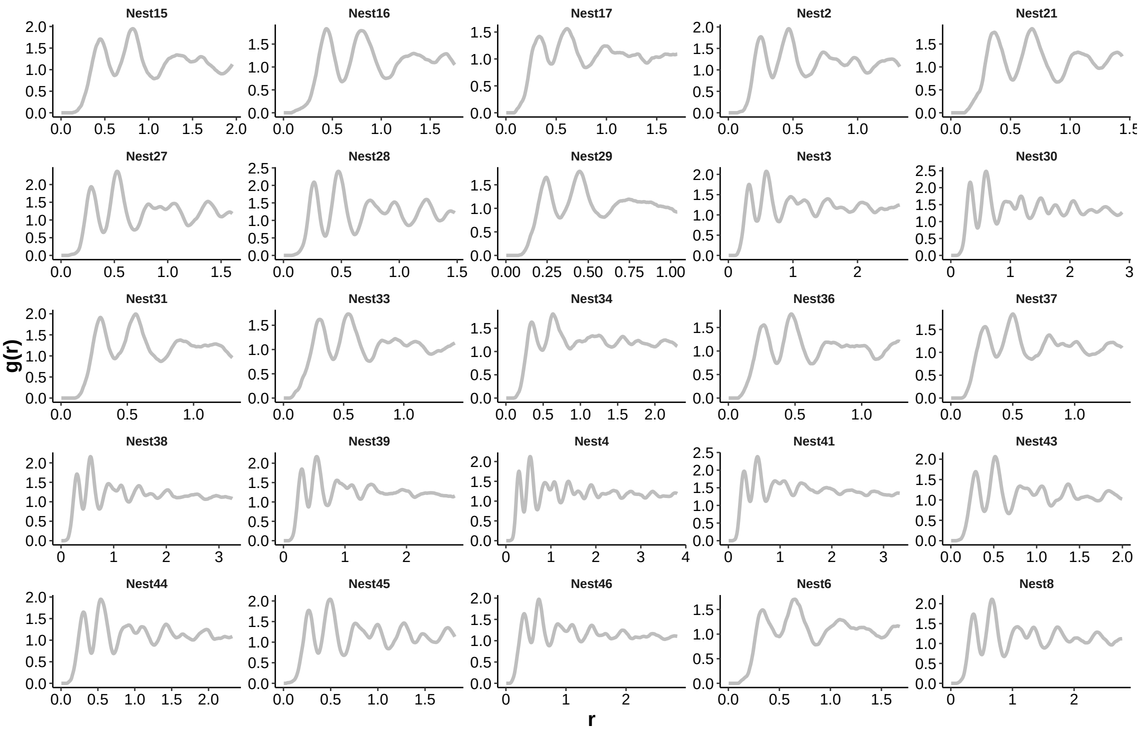


Figure S1: Pair correlation functions of vertex positions for each of the analysed nests. With increasing $r$ (the separation between two vertices,in cm), the curve tends to approach 1 across all the nests.

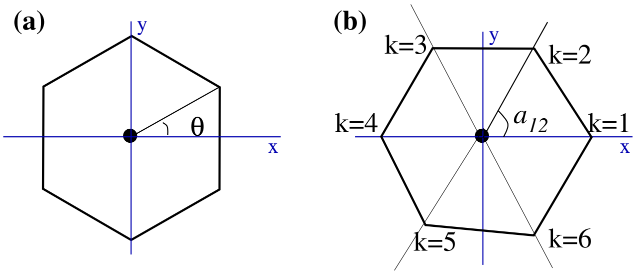


Figure S2: Orientations of a hexagon. (a) A regular hexagon with an orientation angle $\theta$ with respect to the arbitrarily chosen x-axis. For a given $\theta$, $\langle\psi_6\rangle = e^{i6\theta}$, with $0 \le \theta \le \pi/3$. (b) Irregular hexagon constructed by randomly choosing the angles $a_{kl}$.

#### S2. Random orientation—A hexagon with different types of randomness

##### Random Orientations

Consider a regular hexagon as shown in Figure S2(a), with $\theta$ representing the overall orientation with respect to the x-axis. Suppose the orientation angle $\theta$ is random, uniformly distributed in the range$[0, \pi/3)$. The range is dictated by the symmetry that the hexagon returns to an indistinguishable position under a rotation by $\pi/3$. The normalized probability distribution of $\theta$ is,

$$P(\theta) = \frac{3}{\pi}, \quad \text{with} \quad \int_0^{\pi/3} P(\theta)\, d\theta = 1. \tag{S1}$$

For a given $\theta$, the successive angles of a regular hexagon are given by

$$\theta_k = (k-1)\,\frac{\pi}{3} + \theta, \quad k = 1, ...6,$$

so that the cell-orientational order-parameter is

$$\psi_6(\theta) = \frac{1}{6}\sum_k e^{i\theta_k} = e^{i6\theta}.$$

Therefore, the average is given by,

$$\langle\psi_6\rangle = \frac{3}{\pi}\int_0^{\pi/3} e^{i6\theta} d\theta = 0. \tag{S2}$$

This is because for every orientation $\theta < \pi/6$ with $\psi_6 = e^{i6\theta}$, there is an equally probable orientation $(\pi/6)+\theta$ with $\psi_6 = -e^{i6\theta}$, and they cancel each other. In other words, for a disordered situation where a hexagon can take any orientation at random, the average order parameter is zero as mentioned in the text.

#### S3. Random angles

Refer to Figure S2(b).Suppose each of the angles $a_{kk+1}$, $(k = 1, ..., 5)$, subtended by the sides of an irregular hexagon at the cell center is chosen randomly (independent and identically distributed) in the range $(\pi/3) \pm A$ with a probability distribution $P(a) = \frac{1}{2A}$. The value of $A$ is kept small to ensure that $a_{61} > 0$. Let us choose the x-axis such that the vertex numbered 1 is on the axis as in Figure S2(b). The polar angle for site $k$ is,

$$\theta_k = \sum_{j=1}^{k-1} a_{jj+1}, \quad k = 2, ..., 6, \tag{S3}$$

and

$$\langle e^{-i6a} \rangle = \frac{1}{2A} \int_{-A}^{A} e^{i6a}\, da \approx 1 - 6A^2, \quad \text{(for small } A) \tag{S4}$$

Using Eq. S3, we evaluate the average cell-orientational order parameter in terms of $X = \langle e^{-i6a} \rangle$ as,

$$\langle \psi_6 \rangle = \frac{1}{6}\left(1 + X + X^2 + X^3 + X^4 + X^5\right), \tag{S5}$$

which for small $A$ can be written as (from Eq. S4)

$$\langle \psi_6 \rangle \quad \approx \quad \frac{1}{6} \sum_{j=0}^{5} (1 - 6A^2)^j \approx e^{-15A^2}. \tag{S6}$$

Figure S3 shows a comparison of the approximate formula for small $A$ (Eq. (S6)) with numerical simulations. The agreement is good for small $A$.

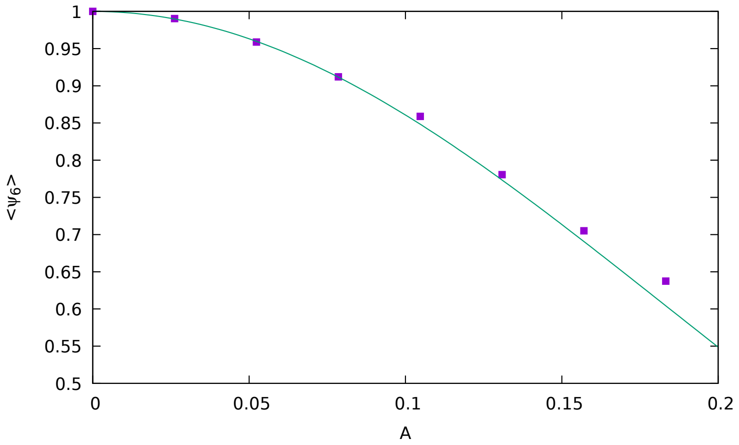


Figure S3: Randomness in irregular hexagon: $\langle \psi_6 \rangle$ vs $A$, where the central angles are uniformly distributed in $(\pi/3 - A, \pi/3 + A)$. Results from numerical simulation using uniform random numbers are shown by the filled squares, while the solid line represents the exponential formula of Eq. (S6).

#### S4. A sample nest

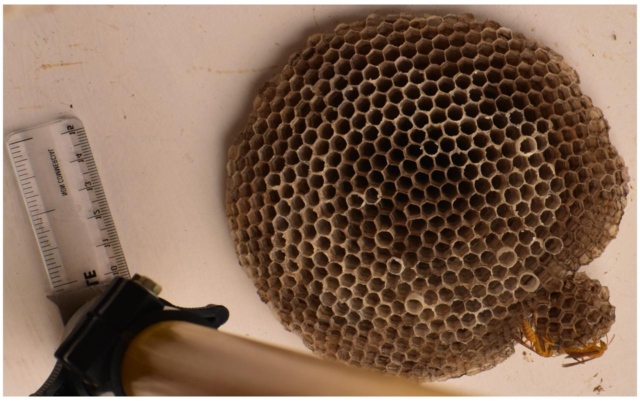

Figure S4: Nest of *Polistes wattii.*

#### S5. On planarity of nests

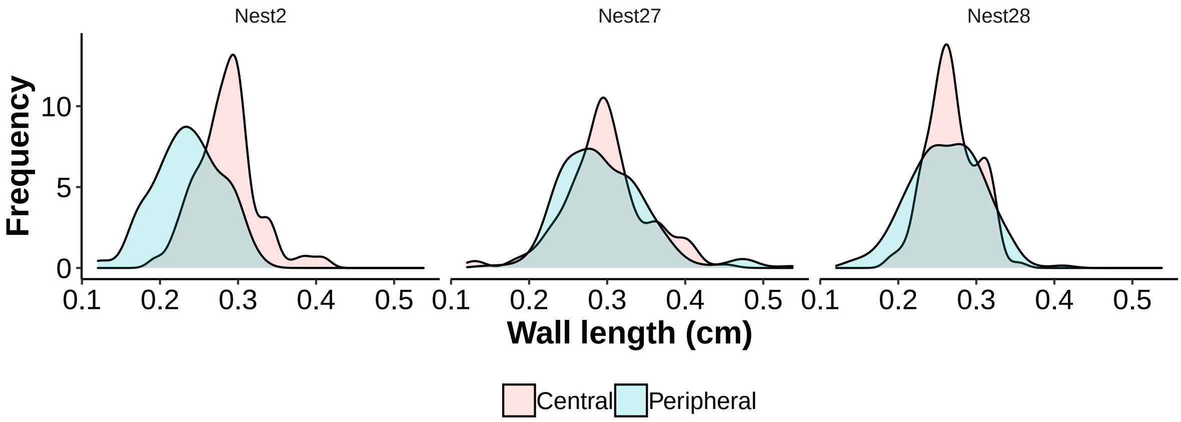


Figure S5: Comparison of wall length distribution between randomly chosen cells that lie in the central and the peripheral regions of three different nests
